# SATORI: A System for Ontology-Guided Visual Exploration of Biomedical Data Repositories

**DOI:** 10.1101/046755

**Authors:** Fritz Lekschas, Nils Gehlenborg

## Abstract

The ever-increasing number of biomedical data sets provides tremendous opportunities for re-use but current data repositories provide limited means of exploration apart from text-based search. Ontological metadata annotations provide context by semantically relating data sets. Visualizing this rich network of relationships can improve the explorability of large data repositories and help researchers find data sets of interest. We developed SATORI—an integrative search and visual exploration interface for the exploration of biomedical data repositories. The design is informed by a requirements analysis through a series of semi-structured interviews. We evaluated the implementation of SATORI in a field study on a real-world data collection.SATORI enables researchers to seamlessly search, browse, and semantically query data repositories via two visualizations that are highly interconnected with a powerful search interface. SATORI is an open-source web application,which is freely available at http://satori.refinery-platform.org and integrated into the Refinery Platform.

## 1 Introduction

Public data repositories are rapidly growing in size and number through implementation of data release policies stipulated by journals and funding agencies (Margolis *et al.*, 2014). The availability of tens of thousands of data sets provides tremendous opportunities for re-use of data across studies. For instance, already published data sets can be used to test a novel hypothesis without having to generate new data.

Alternatively, data from previous studies can be employed as corroborating evidence for observations made in an experiment. Meta studies that include data from dozens or hundreds of published data sets are another common use case for the re-purposing of previously generated data. For example, Lukk *et al.* (2010) studied patterns of gene expression in human tissues based on hundreds of public gene expression data sets and a similar study was conducted for mouse tissues by Zheng-Bradley *et al.* (2010). Other groups have studied connections between different diseases using publicly available data sets (Caldas *et al.*, 2009, 2012; Suthram *et al.*, 2010).

In order to fully embrace sharing and re-use of data, researchers need to be able to i) find data sets, as described by the “*FAIR*” (Wilkinson *et al.*, 2016) principles, and ii) explore data repositories efficiently. As most biomedical raw data is numerical, data sets need to be annotated with metadata to ensure findability. In this context, text-based search is most efficient for navigational queries, e.g., to access a known or recently found data set, and for some transactional queries, e.g., to find the owner of a known data set.

But when the exact context of a data set is unknown or the goal is to learn about the content of a repository, keyword-based search tends to fail (Brandt and Uden, 2003; Holman, 2011) since it provides no overview of the distribution of attributes across the data repository.

Fortunately, an increasing number of data repositories make use of ontologically-annotated metadata, which brings context to annotated data attributes. These semantic relationships can be exploited to relate data sets to each other and to provide an overview at different levels of granularity. To address the needs for precise search and exploration, we propose a system that combines free text and ontologically-annotated metadata. It consists of two interlinked interfaces: a powerful text-based search and a visual analytics exploration tool (Figure 1). In the spirit of Pirolli and Card’s (Pirolli and Card, 2005) sensemaking process, we aim to enable an improved information foraging (Pirolli and Card, 1995) process by enriching search with attribute-based exploration that visualizes the context of data sets and provides a means of semantic top-down exploration, which is a common approach for exploring unknown data or for analyzing collections (Patterson *et al.*, 2001).

Many biomedical data repositories provide a comprehensive text-based search interface, while a few also support other means of exploration. For example, the two major data repositories for gene expression data are Gene Expression Omnibus (GEO) (Barrett *et al.*, 2013) and ArrayExpress (Kolesnikov *et al.*, 2014), each containing over 70,000 data sets as of May 2017. Besides a text-based search, GEO has an indented list facet view for taxonomy groups and provides a list-based repository browser of high-level features such as sample types or organisms. Ar-rayExpress has a combined interface for exploring search results and browsing features via list-based filter options, e.g., organisms, experiment type, or array. Both of these interfaces are insufficient to fully address the needs of scientists for the exploration of published data. Ontology-guided exploration of data repositories also intersects with search visualization and hierarchical data visualization. For example, InfoSky (Andrews *et al.*, 2002) visualizes hierarchical data collections with circular weighted Voronoi treemaps and ResultMaps (Clarkson *et al.*, 2009) groups search results according to a hierarchical classification visualized using the treemap technique. A detailed evaluation of related work is described in Supplementary Section S1^1^. Still,integrative text-based search and ontology-driven visual exploration has not been fully exploited yet.

**Figure 1.**
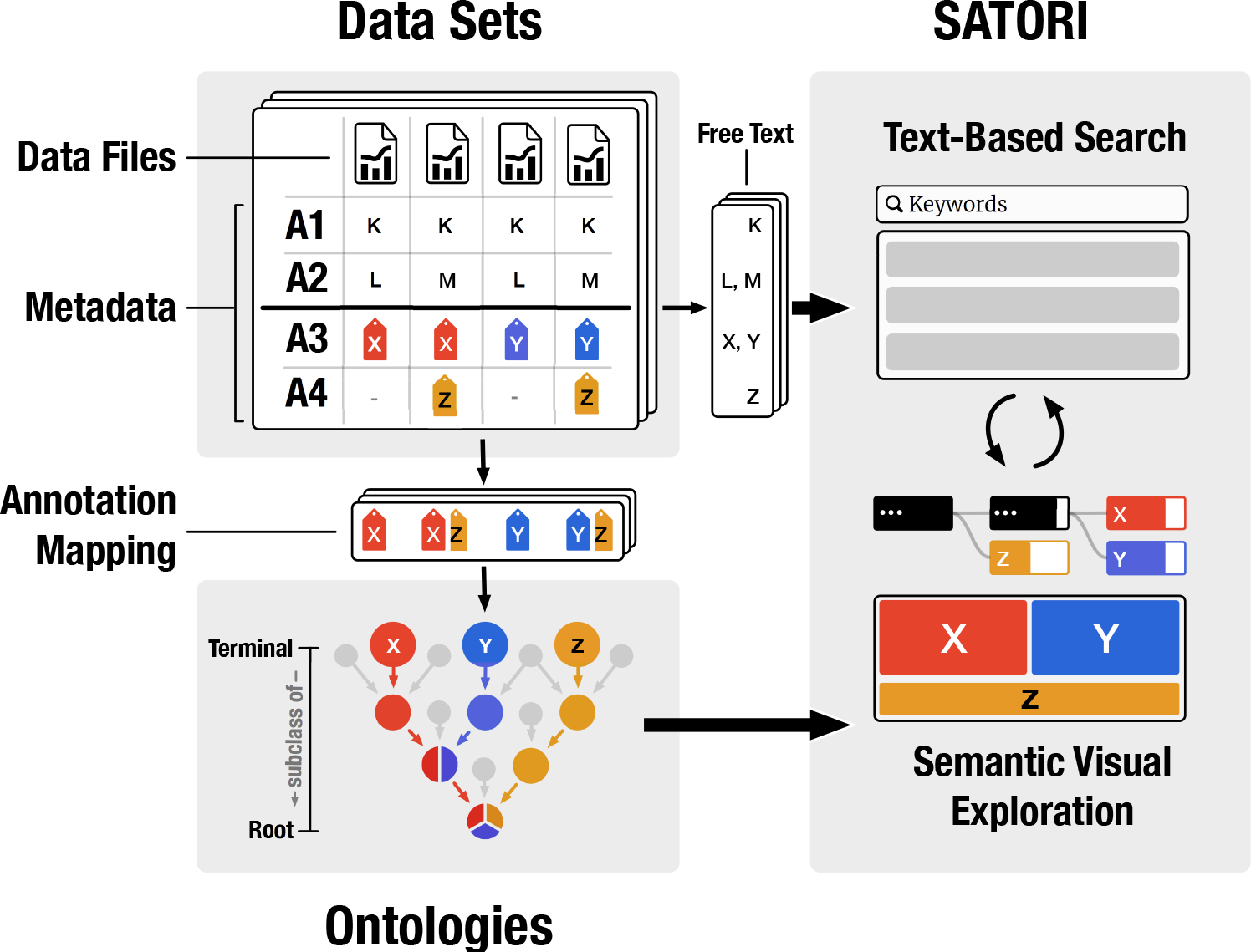
Overview of SATORI’s architecture. Given a collection of data sets, metadata attributes, such as technology, organism, or disease, are extracted and indexed as free text for search and ontologically annotated attribute values (e.g., X, Y, Z) are extracted and linked to their related ontology terms. SATORI combines the unstructured metadata and the structured semantic ontology hierarchy to enhance the explorability of data sets through text-based search and visual exploration.

In this work, we present a novel, ontology-guided visual analytics tool called SATORI (short for **S**emantic **A**nnoTations and **O**ntological **R**elations **I**nterface) that combines search and exploration. First, we identified three user roles, their needs, and the resulting tasks that should be accomplished by any exploration system (Section 2.1) through a series of semi-structured interviews with experts and analyzed the underlying data structure (Section 2.2). Using this as a starting point, we designed two visualizations that represent the content of a data repository. The visualizations provide users with an overview of the attributes used for annotating the data sets and enable semantic querying (Section 3); implementing the foraging loop of the sensemaking process (Pirolli and Card, 2005). We provide a web-based open-source implementation of SATORI for the Refinery Platform^2^ (Section 3.3).

The Refinery Platform is an end-to-end web application for managing, analyzing, and visualizing biomedical data sets. It is build around the Investigation Study Assay (Sansone *et al.*, 2012) data model and relies on a tabular description of data sets. The Refinery Platform provides robust data management and search combined with Galaxy (Afgan *et al.*, 2016) as a powerful analysis back-end. The goal of the Refinery Platform is to integrate the different community standards and therefore needs powerful ways for exploring data sets as a first step in the analysis process.

Using this implementation and two data collections (Stem Cell Commons (Ho Sui *et al.*, 2013) and MetaboLights (Haug *et al*., 2013)), we evaluate our approach in a field study with 6 bioinformaticians and data curators (Section 3.5).

**Figure 2.**
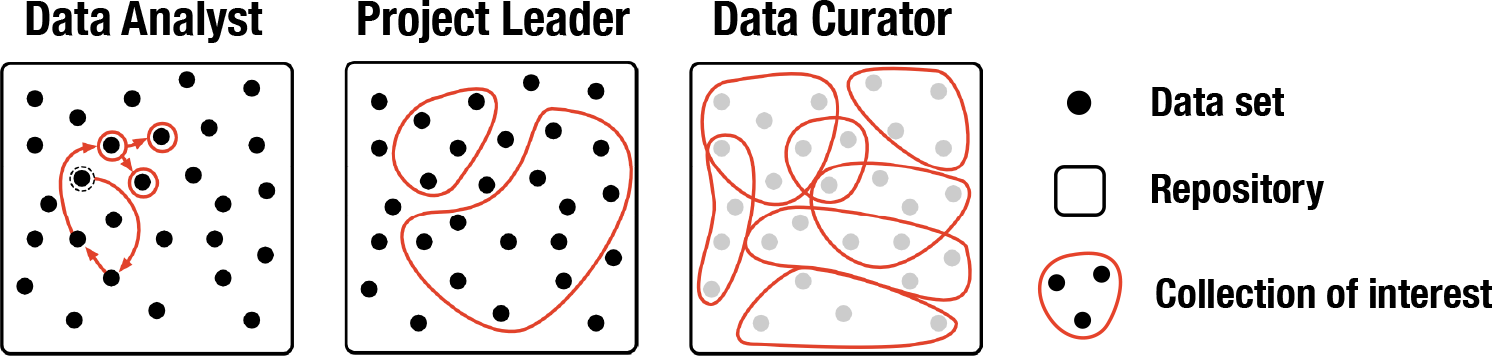
Exploration behavior of different user roles. Data analysts aim at locating specific data sets. Those might be found via *de novo* or iterative search starting from previously known data sets (red arrows). Project leaders focus on collections of data sets and the bigger picture. Data curators are primarily interested in the overall annotation term hierarchy instead of retrieving data sets.

## 2 Methods

### 2.1 Requirements Analysis

We identified three distinct user roles through prior work and user interviews: *data analyst*, *project leader*, and *data curator*. While the roles differ in their needs and tasks, they are not mutually exclusive. To guide our design process, we conducted a requirements analysis to determine the needs and tasks for these roles.

The data analyst is mainly interested in finding relevant data sets that help to answer specific biological questions of interest. Their goal is to analyze data and to subsequently transform it into knowledge. The data analyst might start a *de novo* search or continue from an already known data set to find similar ones (Figure 2). Therefore, the data analyst needs to *(**N1**) find data sets that match specific experimental attributes* and *(**N2**) find data sets that are similar (or dissimilar) to a given collection of data sets*. The project leader focuses on collections of data sets that are potentially important to accomplish a research project. Their goal is to find multiple data sets (Figure 2) and generally investigate whether a data resource is useful. Hence, the project leader needs to *(**N3**) get an overview of the distribution of the experimental attributes across a collection of data sets*. Finally, the data curator is interested in the overall state of curation of the entire data repository and is less concerned about retrieving specific data sets. They need to *(**N4**) get an overview of the annotation term hierarchy and term usage*.

Given the user needs, we derived nine tasks by means of semi-structured interviews with eight PhD-level bioinformaticians (Section S2.3). The first five tasks are related to learning (Marchionini, 2006) about the content of a data repository. First, the user needs to be able to *(**T1**) find the annotation terms of a data set*, i.e., to identify attributes of a data set. To get an idea of the repository’s content the user must also be able to *(**T2**) determine the abundance of annotation terms of a collection of data sets*; e.g., how many data sets are related to *p53*. Furthermore, as the user is often looking for a combination of attributes, it is important to *(**T3**) determine the abundance of multiple annotation terms among a collection of data sets*; e.g., how many data sets are related to *p53*, *liver*, and *RNA-Seq*. Since ontologies provide deep hierarchical descriptions of attributes, it is important to enable the user to *(**T4**) understand annotation term containment relationships*. To aid in determining the relevance of data sets, users must be able to *(**T5**) summarize and view the metadata of data sets*. The next four tasks are related to investigating (Marchionini, 2006) data repositories. First, users must be able to *(**T6**) search for data sets*. It should also be possible to *(**T7**) query by annotation terms*. Knowing the annotation term distribution of a given search, it is also important to be able to *(**T8**) loosen annotation term constraints*. For example, when a search for *human RNA-Seq macrophage* returns insufficient results it might be desirable to include all *monocyte*-related data sets to broaden the results. Finally, *(**T9**) ranking annotation terms* is connected to both learning and interacting. Seeing highly abundant terms can help to get an idea of the main data attributes of the repository. On the other hand, annotation terms with a low abundance can highlight the specifics of some data sets.

A detailed description of the relation between user roles, needs, and tasks is provided in Section S2.

### 2.2 Data

The specifics of data types and structures of biomedical data can vary greatly depending on the research field and application, but the fundamental components for ontology-guided exploration stay the same. As the goal of this work is to find data sets rather than single data files, a data set is regarded as an atomic unit with multiple attributes associated to the files of a data set. Some attributes are linked to ontology terms (called *direct annotation terms*). Given the transitive subclass relationships of ontologies, every superclass of a direct annotation term is also associated to the corresponding data set and denoted as an *indirect annotation term* hereafter. Annotation terms are extracted from the data sets. Hence, the overall number of ontologies and annotation terms depends on the amount to which these data sets have been annotated with ontology terms. More details about the ontology extraction can be found in the Section S4.

**Abstraction**. Ontologies can be considered directed, and in most cases acyclic, graphs in which terms are represented as nodes and relationships as edges between two nodes. For repository exploration, the most important metric of annotation terms is the number of data sets that are associated with to the term. Given a graph G = (*V, E*) with *V* representing the set of vertices and *E* representing the set of edges, we denote the number of times a term *t* has been used to annotate a data set as the *size* of the term. Terms describe sets of data sets; given a term *t*, its set representation is denoted by *S_t_*. The common root ontology term (i.e., *OWL:Thing*) of ontologies imposes an explicit order on the term sets and defines their term hierarchy. The length of the shortest path of a term *t* to the root term is defined as the distance of *t*. In conclusion, the underlying data structure of ontology-annotated data repositories can be described as a semantic polyhierarchy of attributes that describe and organize the data set into groups.

**Figure 3.**
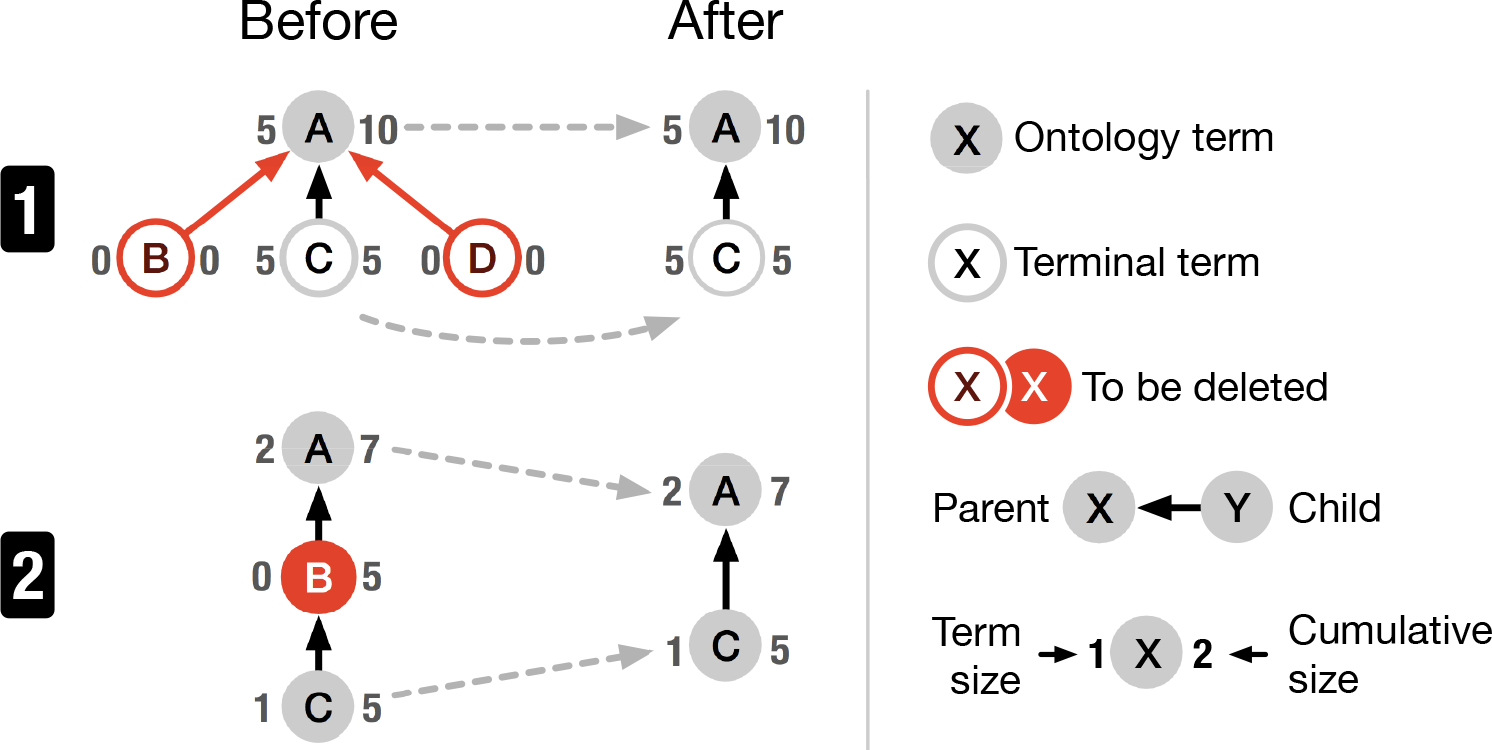
Annotation term pruning: to reduce the size of the annotation term graph and make exploration more efficient, terminal (1) and inner (2) ontology terms of size zero (i.e., terms that have not been used for annotation) are not displayed (red nodes).

**Processing**. Most biomedical ontologies describe a specific domain in its entirety but the number of terms that are used for annotation can be very limited. For example, the Stem Cell Commons (Ho Sui *et al.*, 2013) data collection (Section 3.5) uses only 142 out of 1,269,955 ontology terms. Since the goal of SATORI is to provide a means for finding data sets and understanding the composition of data collections rather than visualizing entire ontologies themselves, unused annotation terms are hidden (Figure 3.1). But even the number of indirectly used terms can be high, given the deep hierarchical structure of some ontologies. Therefore, each parental term should account for a larger collection of data sets than its child term to enable efficient browsing. For example, if a repository contains 10 data sets in total and all data sets are related to *human* then the term *Mammalia* will describe the same 10 data sets as it is an umbrella term that includes human. Hence, the mutual information of all parent terms of *human* related to other attributes (e.g., disease) is zero. Therefore, parental terms of human can be omitted. Thus, the annotation term hierarchy reflects a strict containment set hierarchy. Given three terms *A*, *B* and *C* where *A* is a subclass of *B* and *B* is a subclass of *C*, the set representations *S_A_*, *S_B_* and *S_C_* of the terms should fulfill:

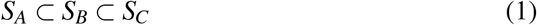

This leads to pruning of terms whose size is zero as illustrated in Figure 3.2. For example, Stem Cell Commons (Ho Sui *et al.*, 2013) includes data sets with files sampled from three species: *human*, *mouse*, and *zebrafish*, which have all been annotated with NCBITaxon (Federhen, 2012). Pruning the subgraph starting from the last common ancestor (*euteleostomi*) according to Equation 1 results in the removal of 37 terms (Figure S3).

## 3 Results

The key goal for SATORI is to enable researchers to more efficiently re-use existing data set by improving the explorability of data repositories through a combination of traditional free-text-based search and context-describing ontology annotations of the metadata.

### 3.1 Interface and Visualizations

SATORI is composed of three interlinked views: data set view, exploration view, and data set summary view. The first two components are visible by default as shown in Figure 4. The data set summary view is displayed on demand (Figure S9) only.

**Data Set View**. The data set view is composed of a text-based search interface (T6), a list of data sets, and a data cart. The design of the search interface has been kept at a minimum to be easy to use Nielsen (1994). A data set is represented by a surrogate—a short description consisting of the title, ownership and sharing information, and an indicator whether the data set is currently saved in the data cart. Additionally, search results feature a *keyword in context* (KWIC) snippet to provide context to the matched keywords (Figure 4.2b and S4). A click on the title of a dataset opens the data set summary view. Being able to quickly get an overview of the metadata of a data set is crucialfor evaluating the relevance of the data set in regards to the information need (T5).

**Figure 4:**
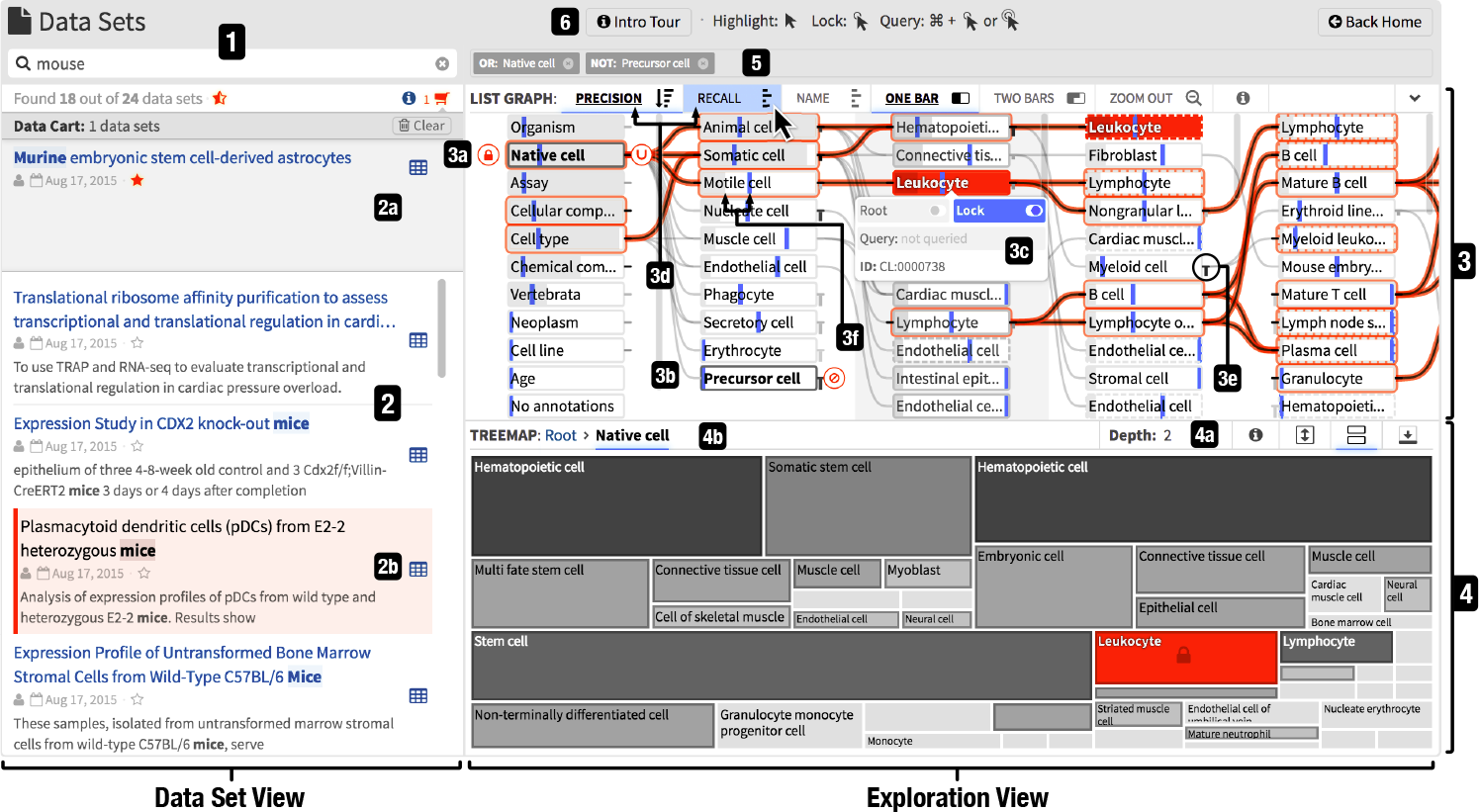
The interface of SATORI consists of the data set and exploration view. The data set view contains the search interface (1) and the list of retrieved data sets (2). Each data set is represented with a surrogate (2b) and can be saved in the data cart (2a). The exploration view is composed of the node-link diagram (3), treemap (4), and query term bar (5). SATORI features several interactive tutorials, indicated by info buttons (6). This example illustrates a query for “native cells” (3a, 4b) excluding “precursor cells” (3b) combined with a synonym keyword search for “mouse” (1); i.e., the retrieved data sets contain the word “mouse” (or synonyms like “mice”) in their free-text description, have been annotated with “native cell”, and do not contain “precursor cell” annotations. Among these retrieved data sets, those annotated with “leukocyte” are highlighted (2b) via the node context menu (3c). The recall, highlighted as blue vertical bars (3d), of leukocyte is less than 50% and the depth of the treemap is set to 2 (4a) to provide a broader overview.

The data cart (Figure 4.2a and S5) integrates into the data set view and enables users to temporarily collect data sets of interest during the exploration process. This reduces the cognitive load during search when comparing results from different searches or annotation term queries as users don’t need to memorize the description of data sets.

**Exploration View**. The exploration view contains two visualizations showing the content of a data repository in terms of the metadata attributes: a node-link diagram (Figure 4.3) and a treemap plot (Figure 4.4). Both display the same data but represent attributes differently to compensate for each other’s limitations. The treemap provides a space-efficient overview of higher-level terms and the node-link diagram represents the relationships between terms across multiple levels.

In the treemap, each term is visualized as a rectangle. The area of the rectangle represents the size of the term relative to its sibling terms. The color indicates the distance to the farthest child term, i.e., the subtree's depth. The farther away a child term is, the darker is the color of the rectangle. The node-link diagram visualizes terms as nodes and links parent and child terms according to subclass relationships defined by the ontologies. Additionally, the node-link diagram shows the precision and recall for each term given the currently retrieved data sets. Precision is defined as the number of data sets annotated with a term divided by the total number of retrieved data sets. Recall is defined as the number of retrieved data sets annotated with a term divided by the total number of data sets annotated with this term across the entire repository. Precision is useful to understand how frequently a term is used for annotation in the retrieved collection of data sets while recall provides a notion of information scent (Pirolli *et al.*, 2000) as it specifies how many relevant data sets for a specific attribute have been retrieved in total (Figure 5).

**Treemap**. The treemap technique visualizes the size of annotation terms (i.e., the number of data sets annotated with a certain term). A main advantage is that the currently selected tree level is always drawn without any overflow or occlusion, which provides an immediate overview. Other visualization techniques that are used for deep hierarchies, e.g., indented lists or node-link diagrams, typically require user interactions to uncover hidden parts. This forces the user to memorize hidden parts. Therefore, the main focus of the treemap is on aggregation of the content by annotation terms. On the other hand, it is hard to perceive the hierarchical structure with treemaps (van Ham and van Wijk, 2002). Also, the rectangle areas are not relative to the root and engender ambiguities (Figure S12). Finally, the area encoding is relatively imprecise compared to other encodings like length (Heer and Bostock, 2010). The node-link diagram compensates for these disadvantages. The treemap also features a breadcrumb path that shows the parental annotation terms to the absolute root term (Figure 4.4b and S8.1) to support T4 and T8. Additionally, the visible depth of the treemap can be increased, via the distance control of the treemap (Figure 4.4a and S8.2). Showing multiple levels of the hierarchy provides better understanding of the structure at the cost of readability.

**Node-Link Diagram**. The node-link technique provides a strong visual notion of connectedness between related annotation terms and emphasizes the hierarchical structure of the annotation term graph (T4). Two terms are visually linked when they are related by an ontological “*is superclass of*” relationship. The directionality follows the reading direction from left to right. To avoid visual clutter, edges are only drawn if both nodes are visible. To indicate the number and position of omitted links, a bar is displayed left or right of a term for incoming and outgoing links to nodes outside the visible area (Figure 4.3e and S10). Annotation terms are ordered in individually sortable (T9) and scrollable columns by their distance to the root term and aligned to the top in order to increase the overall space efficiency (Figure S2). The horizontal layout has been chosen over a vertical layout to make paths follow the reading direction, to easily compare *precision* and *recall* (T2 and T3) within a column via aligned bars, and to provide familiar scroll behavior (i.e., top-down).

By default, the superimposed bar in nodes displays precision and the superimposed vertical line indicates recall. The visual representation for precision and recall depends on which attribute the nodes are ordered by. As users typically start exploring the entire repository, recall is initially equal to 1 and thus less informative than precision. Therefore, by default nodes are sorted by precision. The top navigation bar allows to sort nodes, adjust the bar style, and zoom-out to see the entire graph. In Figure 4.3d nodes are ordered by precision, which is visualized as superimposed gray bars. Recall is highlighted as superimposed blue vertical lines by placing the mouse cursor over the recall button. For example, *motile cell* (Figure 4.3f) has a precision of about 0.2 and a recall of about 0.5.

**Data Set Summary**. The data set summary view supports the *reading and information exporting* step of the information foraging loop (Pirolli and Card, 2005) and addresses T5 (i.e., “*summarize and view the metadata of a data set*”). The layout has been designed to reflect the importance of attributes of data sets for exploration that we derived from the initial semi-structured interviews (Table S2 and S3).

### 3.2 Interactions and Querying

All components of SATORI are linked to provide an integrative exploration experience and to visualize the semantic context of retrieved data sets.

**Identify Data Set-Associated Annotations**. When hovering over a data set surrogate all associated annotation terms are highlighted in the treemap and node-link diagram (T1). Since the node-link diagram shows nodes across different hierarchical levels, direct and indirect annotation terms are handled differently: direct annotation terms are filled in orange while indirect annotation terms only feature an orange outline (Figure 4.3). The node-link diagram can exceed the visible area in size, hence some parts might be occluded. In order to focus on the specific annotation terms of a mouse-hovered data set only, the user can semantically zoom out via a click on the magnifier button such that all terms related to the mouse-hovered data set become visible (Figure S4 and S11).

**Discover Data Sets By Annotations**. To explore the content of a repository based on annotated attribute values, SATORI highlights associated data set surrogates (Figure 4.2b and S6) when moving the mouse cursor over a term in either of the two visualizations (T2). To investigate annotations across views, a click on a rectangle in the treemap or the *Lock* button of the context menu (Figure 4.3c and S7) of the node-link diagram will make the highlighting persistent. Both visualizations support term-based querying to address T7, T8, and T9. A double click on a rectangle in the treemap will zoom into the subtree and simultaneously restrict the retrieved data sets to be associated with the subtree’s root term, hence the data set collection is queried for the clicked term (T7 and T8). The same action can be triggered in the node-link diagram via a click on the *Root* button in the term context menu (Figure 4.3c and S7). Loosening annotation constraints (T9) can be achieved through a click on the treemap’s breadcrumb-like root path view (Figure 4.4b and S8) or by deactivating rooting via another click on the *Root* button in the term context menu. Additionally, the node-link diagram supports more complex Boolean queries for annotation terms with context menu’s *Query* button (T3 and T7). Four query states are implemented: *none*, *or*, *and*, and *not* which the user can toggle through by clicking multiple times on the Query button. Since queries alter the visual state of the visualizations it can get complicated to remember executed queries. Therefore, the query term interface displays all query terms and supports removing queries or altering query states (Figure 4).

**Figure 5.**
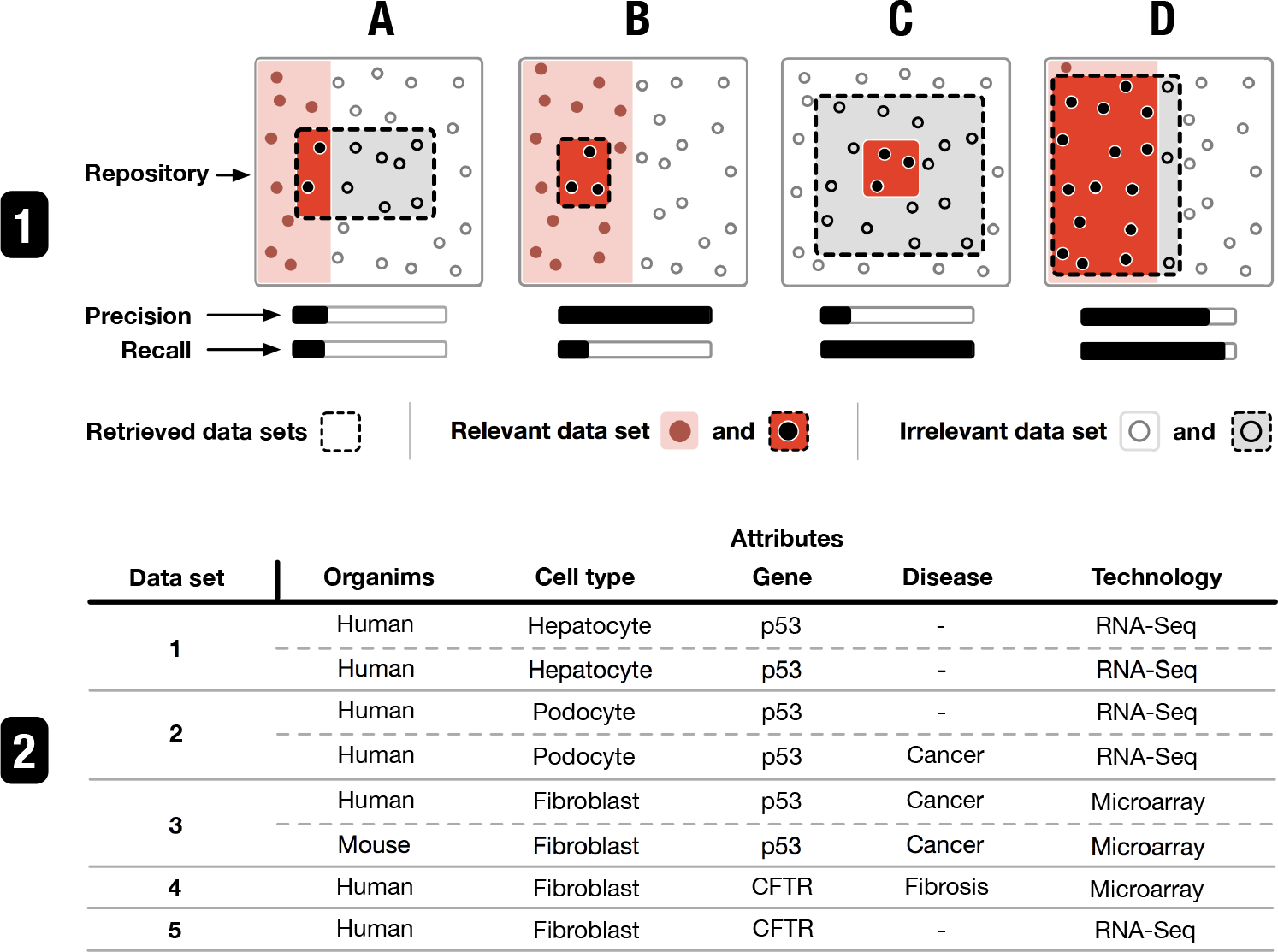
Four cases illustrating the utility of precision and recall. (1a) indicates that an annotation term (i.e., attribute) is not directly related to the search. E.g., given (2), in a broad search for “human”, “fibrosis” is not frequently associated with the data sets found. (1b) states that more data sets related to the annotation term are available. For instance, a search for “human hepatocyte p53” will result in a low recall for *p53* as data sets 2 and 3 are not retrieved since they are not associated with hepatocyte. (1c) illustrates an annotation term that describes a subgroup of all retrieved data sets. E.g., a search for “human fibroblast” shows high recall but low precision for “fibrosis”, indicating that it’s not a commonly studied disease among the retrieved data sets. Finally, (1d) indicates an annotation term that describes many of the retrieved data sets, e.g., a search query for “RNA-Seq” leads to a high recall and precision for p53.

**Figure 6.**
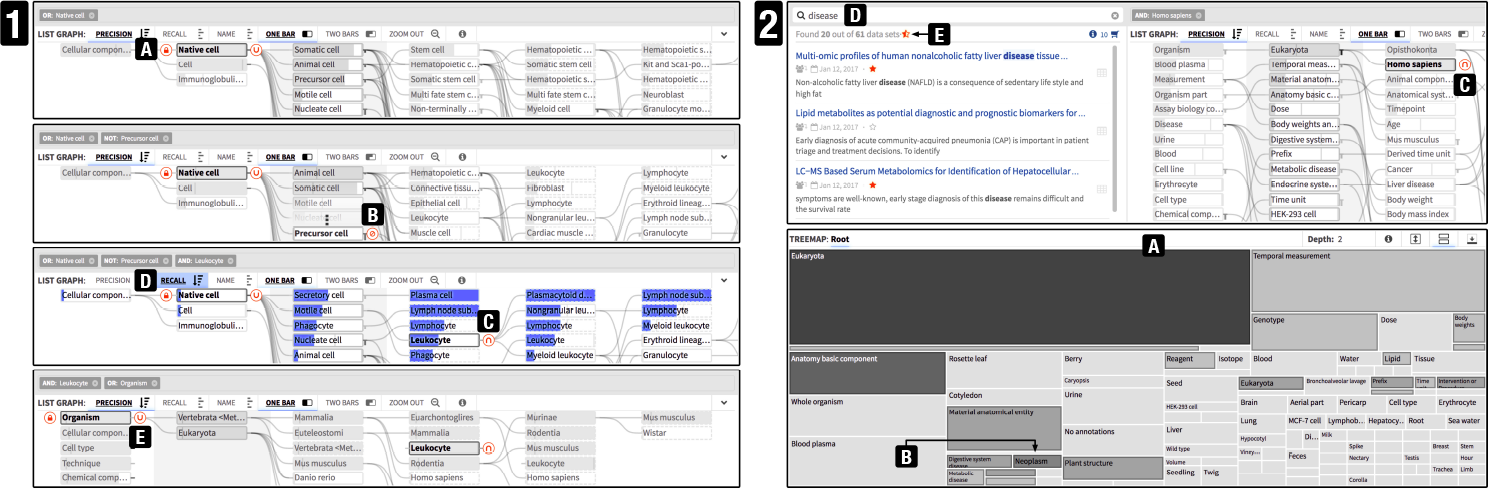
Illustrating steps in two scenarios described in Section 3.4. (1) Four steps in retrieving leukocyte-related data sets, utilizing all four query modes (a, b, and c), recall (d), and inter-branch querying (e). (2) Diverse annotation structure of MetaboLights (a) and the combination of querying and searching (c and d).

### 3.3 Implementation and Scalability

SATORI is a web-based exploration system. The front-end is implemented in JavaScript using D3.js (Bostock *et al.*, 2011) and AngularJS. The information retrieval system is powered by Solr and ontologies are stored in a Neo4J graph database. While Solr provides access to the metadata, Neo4J stores the complete ontology graph. A custom Java plug-in provides access to and retrieves the user specific annotation subgraph for visualization. The Refinery Platform application manages the data set collection and controls the business logic between Solr and Neo4J. Figure S13 shows an overview of SATORI’s architecture. All parts of SATORI are open-source and publicly accessible at GitHub^3^ ^4^, and continuously integrated via Travis-CI to ensure correctness and compatibility.

**Ontology Representation in Neo4J**. We have developed a simplified property graph representation for ontologies to provide space-efficient storage and fast access. Each class is stored as a node with a few core properties like the Uniform Resource Identifier (URI) and label. Superclasses are by default related to the class via a” subclass of“relationship. Other relationships, such as *existential quantification properties*, can be incorporated by adjusting our converter^5^, which translates OWL-formatted ontologies into the property graph model used by Neo4J. To save space when working with multiple ontologies and to speed up node retrieval, every class corresponds to one node, which is labeled with a user-defined ontology abbreviation, e.g., EFO for the Experimental Factor Ontology (Malone *et al.*, 2010). This enables us to traverse a single graph but still trace the occurrence of each class (Section S4).

Since biomedical data repositories can grow quickly, scalability is an important property of any repository exploration system. The performance of SATORI foremost depends on the total number of distinct annotation terms. The impact of the number and size of data sets or ontologies used is negligible since Solr and Neo4J are capable of handling millions of documents and only classes that are directly or indirectly used for annotation are retrieved. Currently, the node-link diagram is the limiting factor as it displays the full annotation graph.We have tested the tool with up to 1000 annotation terms and while the performance decreases, the tool remains usable.

### 3.4 Usage Scenario

In this section we illustrate three usage scenarios for SATORI using the Stem Cell Commons (Ho Sui *et al.*, 2013) and MetaboLights (Haug *et al.*, 2013) data collections, which are comprised of 119 public and a selection of 200 data sets respectively. Data sets of Stem Cell Commons are annotated with 12 ontologies (Table S1) and data sets of MetaboLights are annotated with more than 40 ontologies (Table S2). The Supplementary Video^6^ illustrates the following usage scenarios in detail.

**Context-driven search**. Assume a data analyst is searching for data sets related to leukocytes in Stem Cell Commons. A search query using this term does not retrieve any data sets as no data sets have been explicitly annotated with leukocytes. To evaluate whether similar data sets are available, the data analyst browses the repository by cell types. To restrict the data sets and annotation graph to a specific cell type, the data analyst selects *native cell* as a root term (Figure 6.1a). Three prominent subgroups (*somatic cells*, *animal cells*, and *precursor cells*) become apparent. As leukocytes are not precursor cells the data analyst chooses to filter out data sets related to precursor cells (Figure 6.1b). The next column of child terms contains *leukocyte*, so the data analyst performs an *and* query with it, which retrieves five data sets (Figure 6.1c). Ordering nodes by recall shows that less than 50% of all data sets related to *leukocyte* have been retrieved (Figure 6.1d). The data analyst removes the query terms via the query term bar (Figure 4.5) and retrieves all 12 data sets associated to *leukocytes*. Querying by *leukocyte* and *organism* reveals that the retrieved data sets are associated with *mouse* (Figure 6.1e). Using the data set preview, the analyst finds out that the data sets are related to subtypes of *leukocyte*, which is why the search did not retrieve any data sets.

**State of data curation**. Assume a data curator wants to evaluate the curation state of Stem Cell Commons. The data curator maximizes the treemap visualizations to get an overview of the distribution of the annotation terms. First, the data curator increases the visible depth of the treemap to identify large terminal terms, i.e., rectangles without a border, that represent classes with no subclasses. Such terms could potentially be described in more detail. At depth two, *biotin* pops up as a large terminal node. Zooming into *biotin* reveals that 69 out of 119 public Stem Cell Commons data sets are annotated with it, which is not surprising given that many data sets contain microarray or RNA-Seq experiments. Zooming out to the parent *chemical component* reveals that only one other term, representing a chemical component, is used for annotation, which leaves room for future improvement. Investigating *assay*, another term with two large terminal child terms, reveals that only 116 our of 119 are annotated with *assay*. Using the not query mode, the data curator figures out that the 3 data sets missing the assay annotation are actually related to it as well, in particular *epigenetic modification assay*; highlighting another area for improvement.

**State of data availability**. Assume a project leader wants to learn about the content of Metabolights, in particular diseases-related studies. A first look at the treemap shows that MetaboLights contains a high diversity of studies as represented by many small terminal nodes (Figure 6.2a). The term *neoplasm* pops out given it relative dark background (Figure 6.2b), indicating more specific subtypes. Zooming into *neoplasm* reveals three data sets, which the project leader stores in the data cart for future comparison. The low recall of *disease*, the parent term of *neoplasm*, indicates that more disease related data sets are available. Zooming out and querying for disease using the node-link diagram enables the project leader to check which species have been studied in the context of the disease-related data sets. *Rooting* the node-link diagram for *eukaryotes* reveals that most disease-related studies are associated with *human* and some with *mouse*. Having queried(Figure 6.2c) and saved the 10 *human*-related data sets in the data cart, the project leader clears the annotation query and instead searches for *disease* to find 20 data sets (Figure 6.2d). Some, but not all, of the data sets overlap with the previously-saved data sets (Figure 6.2e). The project leader saves the remaining data sets as well for future project discussions.

### 3.5 Evaluation

We conducted a field study with six bioinformaticians and data curators on the Stem Cell Commons data collection (Section 3.4) to evaluate the general utility of SATORI in exploring a biomedical data repository. Five of the participants are PhD-level scientists and one is a graduate student. The study consisted of a brief introduction to SATORI, a set of *kick-off* tasks for browsing of the Stem Cell Commons data collection (Table S5), and open-ended exploration. During the one-hour-long exploration we recorded anecdotal evidence about SATORI. The participants consist of four data analysts, one project leader, and one data curator to cover all identified user roles (Section 2.1). Three of the six participants were recurring participants from the initial interviews (Section S2.3).

All of the participants stated that SATORI gives them a better understanding of the content of the entire repository compared to a system with only text-based search. Two data analysts mentioned that the ontology-guided exploration interface is very useful for collecting data sets associated with higher-level attribute values, which are not mentioned in the data set description (e.g., *neoplasm* as compared to *glioma*). The project leader mentioned that SATORI significantly aids exploration of unknown big data collections. The data curator said that “*it is really exciting to finally see and explore the (*curated meta*)data*” and that it will be a useful asset for future data curation.

A drawback, identified by all participants, is the difficulty to locate specific terms within the two visualizations. They would like to be able to search for annotation terms. Also, all participants mentioned at the end of the session that SATORI requires some training or introduction. In response, we implemented several step-by-step guides to help first-time users. Also, everyone agreed that many high-level annotation terms are too generic and not useful for exploring data repositories as they are associated with almost all data sets. We have addressed this by defining a set of more useful terms (Table S6 and S7) as entry points. Finally, participants stated that it can currently be time consuming to figure out the current state of querying. We added the query term bar (Figure 4.5), which displays all active queries, to address this concern.

## 4 Discussion

The feedback collected in our field study (Section 3.5) provides evidence that SATORI addresses the needs of the three defined user roles (Section 2.1) and supports the tasks (Section 2.1) in exploring data repositories. Although SATORI is currently integrated into the Refinery Platform, it can be adapted to any data repository that uses controlled and hierarchically-organized vocabularies for annotation. For example, data repositories managed by the European Bioinformatics Institute (EBI) like ArrayExpress (Kolesnikov *et al.*, 2014) or MetaboLights (Haug *et al.*, 2013) already have ontologically-annotated data sets and could easily integrate SATORI into their systems.

A common challenge in visualizing ontology-driven set hierarchies is that many bio-ontologies define complex polyhierarchies with different levels of class granularity. A trade-off has to be made between complexity and usability. Pruning the ontology graph to represent a strict containment hierarchy is a first step but there are more opportunities to improve which class of a pruned branch is kept and to better support visual representation of polyhierarchies. Also, SATORI focuses on exploration of data repositories by finding data sets based on metadata attributes rather than comparing groups of metadata attributes. The ability to compare different groups of annotation terms is currently limited. Existing visualization techniques for exploring set intersections, such as UpSet Lex *et al.* (2014), could be integrated into SATORI to enable richer comprehension of term-related set properties.

## 5 Conclusion

SATORI is a web-based exploration system that combines powerful search with visual browsing to provide an integrated exploration experience. The visualizations serve two purposes: supporting the information foraging loop (Pirolli *et al.*, 2000) and pattern discovery of attribute distributions, as well as ontology-guided semantic querying of the data repository. SATORI contributes to the biomedical domain by unifying text-based search with visual exploration approaches that put data sets into context and shed light in the repository-wide distribution of biological attributes. SATORI extends upon findability of data sets, as described in the FAIR (Wilkinson *et al.*, 2016) principles, by enhancing explorability of data repositories, which is a crucial property as data repositories keep growing at a continuous rate. It is clear that—apart from the design of the visualization and the implementation—the greatest challenge of any semantic exploration approach is that it significantly depends on the quality of data curation. Inconsistent or lacking ontology annotations can result in significant discrepancies between the free-text search and annotation term queries. SATORI also enables curators to evaluate the current state of curation and identify areas that need improvements. We also observed that, due to the nature of the complexities of ontologies, ontology-guided exploration tools require initial learning and are currently most useful for expert users. SATORI focuses on exploration of data repositories given a fixed annotation state. Tracing changes in this annotation space due to ongoing data curation or updated ontologies is an important and unsolved challenge, which requires research on ontology versioning and semantic comparisons.

**Future Work**. To better support locating annotation terms we propose a unified visual query interface, which handles text-based free text search, annotation term search, annotation term query operations, and basic filtering. This would simplify locating specific annotation terms and could encourage more people to explore semantic annotations. While the user study presented in this paper indicates that SATORI is useful for different user types in exploring data repositories, long-term quantitative user studies are needed to evaluate how analysts interact with the system in day-to-day use. Finally, integrating non-ontologically structured metadata into SATORI could have a notable impact as not all descriptive metadata is ontologically annotated.

## Acknowledgements

We would like to thank Burak Alver, Isidro Ciriano, Eunjung Lee, Semin Lee, Youngsook Jung, Jia Wang, Jean Fan, Scott Kallgren, Minseok Kwon, Peter Park, and Shannan Ho Sui for participating in the interviews and the evaluation study to help us design and evaluate SATORI. Furthermore, we would like to express our gratitude to Jennifer Marx, Scott Ouellette, and Ilya Sytchev from the Refinery Platform team, who helped to integrate and deploy SATORI.

## Footnotes

1 Supplementary reference numbers are prefixed with an “S” hereafter.

2 http://refinery-platform.org

3 https://github.com/parklab/refinery-platform

4 https://github.com/flekschas/d3-list-graph

5 https://github.com/flekschas/owl2neo4j

6 https://youtu.be/WpbBoW2f4iM

## References

Afgan, E., Baker, D., Van den Beek, M., Blankenberg, D., Bouvier, D., Čech, M., Chilton, J., Clements, D., Coraor, N., Eberhard, C., et al. (2016). The galaxy platform for accessible, reproducible and collaborative biomedical analyses: 2016 update. Nucleic acids research, 44(W1), W3–W10.

Andrews, K. et al. (2002). The InfoSky Visual Explorer: Exploiting Hierarchical Structure and Document Similarities. Information Visualization, 1(3–4), 166–181.

Barrett, T. et al. (2013). NCBI GEO: archive for functional genomics data sets—update. Nucleic Acids Research, 41(D1), D991–D995.

Bostock, M. et al. (2011). D3: Data-Driven Documents. IEEE Transactions on Visualization and Computer Graphics, 17(12), 2301–2309.

Brandt, D. S. and Uden, L. (2003). Insight into mental models of novice internet searchers. Communications of the ACM, 46(7), 133–136.

Caldas, J. et al. (2009). Probabilistic retrieval and visualization of biologically relevant microarray experiments. Bioinformatics, 25(12), i145–i153.

Caldas, J. et al. (2012). Data-driven information retrieval in heterogeneous collections of transcriptomics data links SIM2s to malignant pleural mesothelioma. Bioinformatics (Oxford, England), 28(2), 246–253.

Clarkson, E. et al. (2009). ResultMaps: Visualization for Search Interfaces. IEEE Transactions on Visualization and Computer Graphics, 15(6), 1057–1064.

Federhen, S. (2012). The ncbi taxonomy database. Nucleic acids research, 40(D1), D136–D143.

Haug, K. et al. (2013). MetaboLights—an open-access general-purpose repository for metabolomics studies and associated meta-data. Nucleic Acids Research, 41(D1), D781–D786.

Heer, J. and Bostock, M. (2010). Crowdsourcing graphical perception: using mechanical turk to assess visualization design. In Proceedings of the SIGCHI Conference on Human Factors in Computing Systems, CHI’ 10, pages 203–212. ACM.

Ho Sui, S. et al. (2013). The Stem Cell Commons: an exemplar for data integration in the biomedical domain driven by the ISA framework. AMIA Joint Summits on Translational Science proceedings AMIA Summit on Translational Science, 2013, 70.

Holman, L. (2011). Millennial students’ mental models of search: Implications for academic librarians and database developers. The Journal of Academic Librarianship, 37(1), 19–27.

Kolesnikov, N. et al. (2014). ArrayExpress update—simplifying data submissions. Nucleic Acids Research, page gku1057.

Lex, A. et al. (2014). UpSet: Visualization of Intersecting Sets. IEEE Transactions on Visualization and Computer Graphics (InfoVis’14), 20(12), 1983–1992.

Lukk, M. et al. (2010). A global map of human gene expression. Nature Biotechnology, 28(4), 322–324.

Malone, J., Holloway, E., Adamusiak, T., Kapushesky, M., Zheng, J., Kolesnikov, N., Zhukova, A., Brazma, A., and Parkinson, H. (2010). Modeling sample variables with an experimental factor ontology. Bioinformatics, 26(8), 1112–1118.

Marchionini, G. (2006). Exploratory search: from finding to understanding. Communications of the ACM, 49(4), 41–46.

Margolis, R. et al. (2014). The national institutes of health’s big data to knowledge (bd2k) initiative: capitalizing on biomedical big data. Journal of the American Medical Informatics Association, 21(6), 957–958.

Nielsen, J. (1994). Usability Engineering. Elsevier.

Patterson, E. S. et al. (2001). Predicting Vulnerabilities in Computer-Supported Inferential Analysis under Data Overload. Cognition, Technology & Work, 3(4), 224–237.

Pirolli, P. and Card, S. (1995). Information foraging in information access environments. In Proceedings of the SIGCHI conference on Human factors in computing systems, pages 51–58. ACM Press/Addison-Wesley Publishing Co. bibtex: pirolli_information_1995-1.

Pirolli, P. and Card, S. (2005). The sensemaking process and leverage points for analyst technology as identified through cognitive task analysis. In Proceedings of International Conference on Intelligence Analysis.

Pirolli, P. et al. (2000). The Effect of Information Scent on Searching Information: Visualizations of Large Tree Structures. In Proceedings of the Working Conference on Advanced Visual Interfaces, AVI’00, pages 161–172, New York, NY, USA. ACM.

Sansone, S.-A., Rocca-Serra, P., Field, D., Maguire, E., Taylor, C., Hofmann, O., Fang, H., Neumann, S., Tong, W., Amaral-Zettler, L., et al. (2012). Toward interoperable bioscience data. Nature genetics, 44(2), 121–126.

Suthram, S. et al. (2010). Network-Based Elucidation of Human Disease Similarities Reveals Common Functional Modules Enriched for Pluripotent Drug Targets. PLoS Computational Biology, 6(2).

van Ham, F. and van Wijk, J. J. (2002). Beamtrees: compactvisualizationoflargehierarchies. In IEEE Symposium on Information Visualization, 2002. INFOVIS 2002, pages 93–100. IEEE.

Wilkinson, M. D. et al. (2016). The fair guiding principles for scientific data management and stewardship. Scientific data, 3.

Zheng-Bradley, X. et al. (2010). Large scale comparison of global gene expression patterns in human and mouse. Genome Biology, 11(12), 1–11.

